# Altered Thalamocortical Connectivity in Six-Week Old Infants at High Familial Risk for Autism Spectrum Disorder

**DOI:** 10.1101/2020.06.07.139147

**Authors:** Aarti Nair, Rhideeta Jalal, Janelle Liu, Tawny Tsang, Nicole M. McDonald, Lisa Jackson, Carolyn Ponting, Shafali S. Jeste, Susan Y. Bookheimer, Mirella Dapretto

## Abstract

Converging evidence from neuroimaging studies has revealed altered connectivity in cortical-subcortical networks in youth and adults with autism spectrum disorder (ASD). Comparatively little is known about the development of cortical-subcortical connectivity in infancy, before the emergence of overt ASD symptomatology. Here we examined early functional and structural connectivity of thalamocortical networks in infants at high familial risk for ASD (HR) and lowrisk controls (LR). Resting-state functional connectivity (rs-fcMRI) and diffusion tensor imaging (DTI) data were acquired 52 six-week-old infants. Functional connectivity was examined between six cortical seeds –prefrontal, motor, somatosensory, temporal, parietal, and occipital regions– and bilateral thalamus. We found significant thalamic-prefrontal underconnectivity, as well as thalamic-occipital and thalamic-motor overconnectivity in HR infants, relative to LR infants. Subsequent structural connectivity analyses also revealed atypical white matter integrity in thalamic-occipital tracts in HR infants, compared to LR infants. Notably, aberrant rs-fcMRI and DTI connectivity indices at 6 weeks predicted atypical social development between 6 and 36 months of age, as assessed with eye-tracking and diagnostic measures. These findings indicate that thalamocortical connectivity is disrupted at both the functional and structural level in HR infants as early as six weeks of age, providing a possible early marker of risk for ASD.

## INTRODUCTION

Autism spectrum disorder (ASD) is a highly heritable neurodevelopmental condition associated with altered brain development (Ecker 2017). While the etiology of ASD remains poorly understood (Chen et al. 2015; Lyall et al. 2017; Constantino 2018), a large body of neuroimaging studies have found abundant evidence implicating both functional and structural connectivity abnormalities in ASD (Ecker et al. 2015; Hernandez et al. 2015; Rane et al. 2015). Resting-state functional connectivity (rs-fcMRI) studies have suggested that ASD is characterized by disrupted connectivity across several brain regions, including atypical connectivity within the default mode (Uddin, Supekar and Menon 2013; Washington et al. 2014; Hull et al. 2017; Padmanabhan et al. 2017), salience (Uddin, Supekar, Lynch, et al. 2013; Abbott et al. 2016; Green et al. 2016; Margolis et al. 2019), central executive (Abbott *et al.* 2016; Menon 2018; Lawrence et al. 2019), attention (Farrant and Uddin 2016; Duan et al. 2017), social cognition (von dem Hagen et al. 2013; Olivito et al. 2017; Chen et al. 2018; Muller and Fishman 2018), visual (Keown et al. 2013; Chen *et al.* 2018), and sensorimotor networks (Nebel et al. 2014; Cerliani et al. 2015; Green et al. 2017). Collectively, these studies have characterized such atypicalities in functional connectivity as predominantly reflecting underconnectivity within regions involved in higher-order cognitive and social functions, coupled with overconnectivity within primary sensory brain areas.

Additionally, a growing body of work using diffusion tensor imaging (DTI) has also identified atypical structural connectivity in youth with ASD (Di et al. 2018). Indeed, white matter tract integrity has been found to be compromised in individuals with ASD within the corpus callosum, cingulum, projections fibers within subcortical regions (internal/external capsule, anterior thalamic radiation), cerebellum, and associations fibers within the frontal, parietal, temporal and occipital lobes such as the uncinate fasciculus, arcuate fasciculus, superior longitudinal fasciculus, inferior longitudinal fasciculus, and fronto-occipital fasciculus in children and adults with ASD (Rane et al. 2015; Ismail et al. 2016; Yamasaki et al. 2017). Recent studies have also shown altered white matter integrity in females with ASD relative to neurotypical females (Rane *et al.* 2015). Despite little consistency in the localization of the reported atypicalities across studies, the existing literature support the hypothesis of abnormal development of both functional networks and white matter tracts in ASD.

Neuroimaging studies to date have largely focused on children and adolescents or adults with ASD. While ASD symptomatology typically emerges in the first years of life, there have been comparatively fewer neuroimaging studies with toddlers and younger children with ASD, likely due to the challenges in conducting MRI studies in this age group. In recent years, however, several neuroimaging studies have been undertaken in infants at high familial risk for ASD (i.e., siblings of children with ASD) who have a much higher occurrence rate than the general population (Ozonoff et al. 2011). Prospectively following these high-risk (HR) infants provides the unique opportunity to study the neural antecedents of risk for developing ASD prior to the onset of overt symptoms. Thus, examining brain connectivity in these young HR infants may help identify early biomarkers of ASD and, relatedly, the deployment of earlier and more effective interventions. Many studies in HR infants have focused on structural brain development; this work has identified enlarged postnatal brain volume (including greater grey matter, white matter, as well as cerebellar and subcortical volume; Hazlett et al. 2017; Swanson et al. 2017; Wolff et al. 2018; Pote et al. 2019), and increased extra-axial fluid (Shen et al. 2017; Shen and Piven 2017) starting as early as six months of age. Studies using DTI in HR infants have identified alterations in white matter tracts connecting frontal, temporal, and subcortical regions, as well as reduced local and global efficiency in occipital, parietal, and temporal lobe tracts between ages 6-36 months (Lewis et al. 2014; Conti et al. 2015; Lewis et al. 2017; Shen and Piven 2017; Wolff *et al.* 2018). Few studies have also found that white matter tract trajectories have a unique pattern of increased FA at age six months followed by slower subsequent growth in HR infants who later developed ASD compared to their non-ASD peers (Wolff et al. 2012; Solso et al. 2016). Interestingly, atypical lateralization of dorsal language tracts has also been observed in HR infants as young as 6 weeks of age, which predicted later language development as well as ASD symptomatology (Liu et al. 2019).

In recent years, studies using rs-fcMRI in HR infants have also began to emerge linking early alterations in functional connectivity to subsequent ASD symptomatology. Notably, (Emerson et al. 2017) found that whole-brain rs-fcMRI metrics obtained at six months of age and their association with behavioral measures could distinguish between HR infants who were later diagnosed with ASD at age 24 months from those who did not. Other rs-fcMRI studies have implicated specific networks. For instance, atypical functional connectivity in the default mode, dorsal attention, and visual networks has been linked to poor initiation of social attention in HR infants between ages 12-24 months (Eggebrecht et al. 2017). In addition, altered patterns of connectivity in these networks, as well as in control and subcortical networks, have been shown to be associated with stereotyped and restricted behaviors in HR infants between 12 and 24 months of age (McKinnon et al. 2019). Of note, two recent rs-fcMRI studies reported atypical development of language-related networks from 1.5 to 9 months of age in HR infants (Liu et al., in Press), as well as altered functional connectivity in social brain regions in HR newborns, suggesting that the neural underpinnings of atypical language and social development in HR infants may be detectable at birth, or in the first few weeks of life.

In parallel to the neuroimaging studies described above, recent work using eye-tracking methods to assess social visual engagement in HR infants has greatly contributed to the search for early neuroendophenotypes of ASD (Klin, Schultz & Jones, 2015). HR infants have been shown to exhibit atypical gaze patterns while viewing social stimuli in the first year of life (Elsabbagh et al. 2012; Chawarska et al. 2013; Elsabbagh et al. 2013), as well as enhanced visual search abilities and a preference for geometric figures over social scenarios (Gliga et al. 2015; Pierce et al. 2016). Importantly, this aberrant processing of socially-relevant information as measured by eye-tracking techniques has been found to be strongly influenced by genetic factors as well as to predict later diagnosis of ASD (Jones et al 2013; Constantino, 2017). To our knowledge, only one study to date has examined the neural correlates of social-visual engagement in HR infants combining both neuroimaging and eye-tracking techniques (Tsang under review). Interestingly, greater connectivity between salience network hubs and prefrontal regions at six weeks of age was associated with increased attention to faces in the first year of life only; in contrast, higher salience network connectivity with sensorimotor regions in HR infants was associated with later atypicalities in social and sensorimotor development. Similar to prior functional imaging studies in HR infants, this study focused primarily on cortical network connectivity patterns. However, theoretical models have posited a switch from more reflexive, externally-driven sensory responses under subcortical control to experience-dependent responses (e.g., social engagement) under cortical control around 6-8 weeks of age (Johnson and Mareschal 2001; Klin, Shultz, et al. 2015; Shultz et al. 2018). Hence, examining cortical-subcortical connectivity and its relationships to behavioral phenotypes of early social development during this crucial age window is essential to foster our understanding of the pathogenesis of ASD risk.

Of particular interest is the thalamus, a critical subcortical structure involved in the integration of visual, auditory, and somatomotor functions. During development, thalamocortical pathways provide crucial input for the specialization of the neocortex (Schlaggar and O’Leary 1991; Stojic et al. 1998; O’Leary and Nakagawa 2002). Normative patterns of functional thalamocortical connectivity have been identified in neurotypical individuals using rs-fcMRI (Zhang et al. 2008; Fair et al. 2010; Zhang et al. 2010). Studies on children and adolescents with ASD have demonstrated both reduced functional thalamocortical connectivity, especially for thalamic-prefrontal networks, and overconnectivity within thalamic-temporal networks (Nair et al. 2013; Nair et al. 2015). In addition to reports of aberrant functional thalamocortical connectivity, there is also evidence of alterations in structural thalamocortical connectivity in ASD, with higher mean (MD) and radial (RD) diffusivity indices for motor and somatosensory tracts in individuals with ASD as compared to neurotypical peers (Nair *et al.* 2013). In another study by (Nair *et al.* 2015), diffusivity indices between thalamus and several prefrontal, parietal, and temporal regions were also impacted with reduced fractional anisotropy (FA) within thalamic-prefrontal tracts and higher MD, RD, and axial (AD) diffusivity within tracts connecting the thalamus and these regions. Notably, these aberrant patterns of functional and structural thalamocortical connectivity in ASD individuals were found to be associated with poor social interaction skills, executive functioning difficulties, restricted and repetitive behaviors, and sensorimotor atypicalities.

Despite the critical role of the thalamus for the early specialization of the neocortex and evidence implicating altered thalamocortical connectivity in ASD symptomatology (Schlaggar and O’Leary 1991; Stojic *et al.* 1998; O’Leary and Nakagawa 2002; Nair *et al.* 2013; Nair *et al.* 2015; Chen et al. 2016; Green *et al.* 2017; Woodward et al. 2017; Linke et al. 2018; Iidaka et al. 2019), relevant evidence about the thalamus and its functional and structural connections in HR infants remains limited. Only one study thus far has examined the relationship between structural measures of thalamic functioning and both verbal and non-verbal skill acquisition between 6 and 24 months of age in HR infants, reporting that atypical patterns of language development were associated with thalamic volume in this group (Swanson *et al.* 2017). Thus, our objective for the current study was to examine both functional and structural thalamocortical connectivity in HR infants during this pivotal developmental transition and to examine whether early atypicalities would predict risk for later developing overt symptoms of ASD. Our hypotheses were that both functional and structural thalamocortical networks would be impacted in HR infants as early as 6 weeks of age. Similarly to what previously observed in older children and adolescents with ASD (Nair *et al.* 2013; Nair *et al.* 2015), we expected underconnectivity within thalamic-prefrontal networks and overconnectivity with thalamic-temporal networks along with accompanying atypical diffusivity indices within these networks. We also predicted that these network alterations would be associated with later behavioral phenotypes of ASD such as decreased preferential viewing of social stimuli using eye-tracking, and social interaction deficits on diagnostic measures in the first three years of life.

## METHODS

### Participants

Infants were recruited as part of a longitudinal developmental project at University of California, Los Angeles (NIH ACE-II) examining early signs of ASD risk from birth to age 36 months. Our sample consisted of 52 6-week-old infants who successfully underwent a rs-fcMRI scan during natural sleep, including 24 HR infant siblings of children diagnosed with ASD and 28 LR infants. DTI data were also collected for a subset of 47 infants, (HR=24 HR; LR=23); of whom 17 HR and 15 LR infants for the DTI analyses (see DTI processing section below). See Table 1 for demographic details of each groups. Exclusionary criteria for both groups included prematurity (birth before 35 weeks of gestation), as well as serious medical and neurological conditions (i.e., perinatal stroke, hydrocephalus). Age was corrected to 40 weeks of gestation for infants born prior to 38 weeks. Written informed consent was obtained from all participants’ caregivers. The study protocol was approved by the UCLA Institutional Review Board.

**Table 1.** Demographic information for HR and LR groups for rs-fcMRI and DTI data

|  | **Rs-fcMRI** |  |  | **DTI** |  |  |
| --- | --- | --- | --- | --- | --- | --- |
|  | **LR**<br>**N=28** | **HR**<br>**N=24** | **P** | **LR**<br>**N=15** | **HR**<br>**N=17** | **P** |
| Age at Scan<br>(Weeks) | 6.65±(1.20) | 6.34±(1.21) | .22 | 6.43±(1.27) | 6.42±(1.08) | .96 |
| Gestation Length<br>(Weeks) | 37.99±(0.29) | 37.70±(0.82) | .52 | 38.82±(1.51) | 39.73±(0.88) | .07 |
| *Sex* |  |  | .21 |  |  | .31 |
| Female | 12 (43%) | 11 (46%) |  | 4 (27%) | 7 (41%) |  |
| Male | 16 (57%) | 13 (54%) |  | 11 (73%) | 10 (59%) |  |
| *Race* |  |  | .43 |  |  | .36 |
| White | 22 (79%) | 13 (54%) |  | 12 (80%) | 10 (59%) |  |
| Non-white | 6 (21%) | 11 (46%) |  | 3 (20%) | 7 (41%) |  |
| *Family Income* |  |  | .18 |  |  | .82 |
| Not Answered | 1 (4%) | 2 (8%) |  | 0 | 2 (12%) |  |
| <50 K | 3 (11%) | 5 (21%) |  | 2 (13%) | 3 (18%) |  |
| 50-75 K | 5 (18%) | 4 (17%) |  | 2 (13%) | 2 (12%) |  |
| 75-100 K | 6 (21%) | 5 (21%) |  | 4 (27%) | 4 (23%) |  |
| 100-125 K | 4 (14%) | 2 (8%) |  | 2 (13%) | 2 (12%) |  |
| >125 K | 9 (32%) | 6 (25%) |  | 5 (33%) | 4 (23%) |  |

### Imaging Data acquisition

Participants underwent MRI at age 6 weeks on a 3T Siemens Trio scanner using a 12-channel head coil during natural sleep. Parents were instructed to put their infant to sleep using their typical routine before transferring the infant to the scanner bed. Infants were placed on a custom-made bed situated inside the head coil and secured with a Velcro strap. Pliable and soft silicone earplugs and MiniMuffs Neonatal Noise Attentuators (Natus Medical Inc., San Carlos, CA) were used as hearing protection. A weighted blanket and foam pads were used to minimize motion. Additionally, a research staff member remained in the scanner room during the scan to monitor infants for waking, movement, or signs of distress. A scout localizer was used for alignment. The rs-fcMRI scan was acquired using an EPI gradient-echo acquisition lasting 8 minutes, covering the whole cerebral volume (TR=2000ms, TE=28ms, FOV=192mm, 34 slices, voxel size=3×3×4mm). A high-resolution structural T2-weighted echo-planar imaging volume (spin-echo, TR=5000ms, TE=34ms, FOV=192mm, 34 slices, voxel size=1.5×1.5×4mm) was acquired coplanar with the functional scans to ensure identical distortion characteristics of the images. The DTI scans (TR=9500ms, TE=87ms, FOV=256mm, 75 slices, voxel size=2×2×2mm) consisted of 32 diffusion-weighted directions (b=1000), three scans without any diffusion sensitization (b=0), and six additional directions (b=50).

### Behavioral Measures

To examining the relationship between thalamocortical networks and early social correlates of risk for ASD, we examined behavioral measures that were collected across multiple timepoints as part of the broader longitudinal project protocol. These measures included a free-view eye-tracking task of animated social interaction, and diagnostic assessments of ASD risk including clinician administered measures and caregiver reports.

For the eye-tracking task, video stimuli were a 2-min segment of ‘A Charlie Brown Christmas’ administered at ages 6, 9, and 12 months of age using a Tobii T60XL system (Tsang et al. 2019). This video stimuli (See Supplementary Figure 1) consisted of animated characters conversing (~ 80 s), cheering and dancing in a group (~20 s), and speaking to a group of characters (~20 s). Caregivers held their infant on their lap for the duration of the video, approximately 60cm from the computer monitor (65-cm screen; 720×480 pixels resolution; <.05° visual angle; 60hz temporal resolution). A 9-point calibration scheme was repeatedly used prior to data collection until the infant’s point of gaze (PoG) was within 1° of the center of the target. The video stimuli were presented only after the calibration criterion had been reached to ensure accuracy in PoG data to be included in the analyses.

Diagnostic measures of risk for ASD symptoms were administered at different time points between 12 and 36 months of age. At 12 months, we completed the Autism Observation Scale for Infants (AOSI; Bryson et al. 2008) – an 18-item observational measure used to identify and track early signs of ASD administered by expert clinicians. At 24 months, caregiver ratings were obtained on the Social Responsiveness Scale, Second Edition – Preschool version (SRS-P; Constantino 2012) – a 65-item checklist of ASD symptomatology with ratings ranging from 0 (not true) to 3 (almost always true). Finally, at 36 months, the Autism Diagnostic Observation Schedule, Second Edition (ADOS-2; Lord 2012) – a semi-structured standardized assessment of ASD symptoms (meeting formal diagnostic criteria) was administered by expert clinicians who were research-reliable on this measure.

### Data Analysis

#### Regions of interest

The UNC Infant 0-1-2 neonate atlas (Shi et al. 2011) was used to obtain bilateral masks for six cortical seeds: motor, occipital, parietal, prefrontal, somatosensory, and temporal regions (Figure 1A). The masks for these cortical regions of interest (ROIs) were created by combining anatomical parcellations from this atlas (see Supplementary Table 1 for a complete listing of parcellations comprising each cortical ROI). For DTI analyses, white matter tracts connecting only those thalamocortical regions which demonstrated impacted functional connectivity findings were examined.

**Figure 1.**
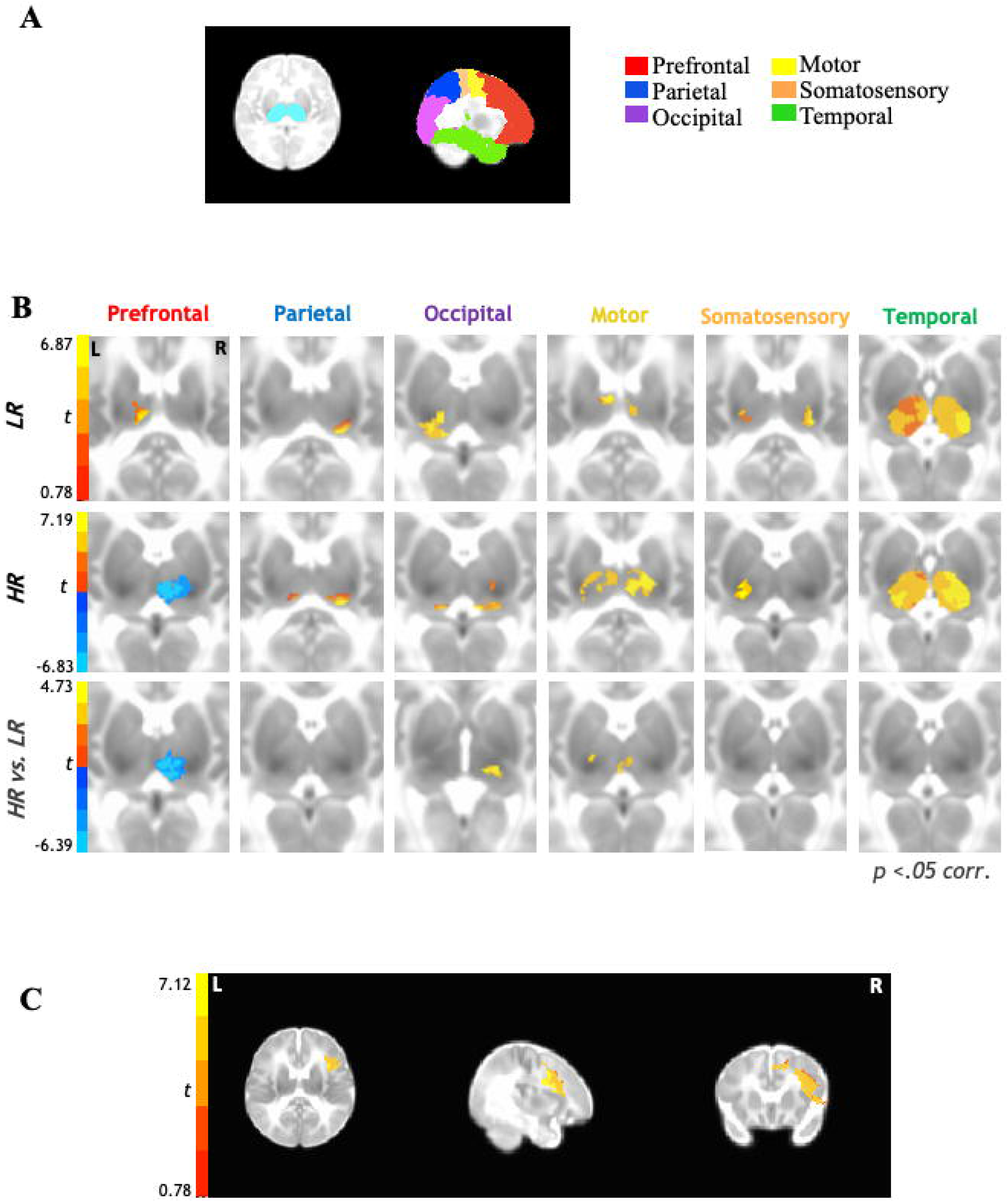
Functional thalamocortical connectivity results for 6-week old infants at higher-risk (HR) for developing ASD compared to their low-risk (LR) peers. **(A.)** Bilateral thalamic masks and cortical seeds of interest. **(B.)** Partial correlation analysis results indicate thalamocortical connectivity for the LR group (top row) consistent with prior literature with typically-developing children and adults. In contrast, HR group shows marked underconnectivity for prefrontal-thalamic networks (middle row). Between-samples paired t-test results indicate that HR group shows additional overconnectivity effects for occipital-thalamic and motor-thalamic networks as compared to LR group (bottom row). **(C.)** Reverse connectivity analysis using thalamic clusters of underconnectivity with prefrontal ROI for HR group implicates that the right inferior frontal gyrus as driving the prefrontal-thalamic underconnectivity findings.

#### Rs-fcMRI Data Processing

Functional images were processed using the Analysis of Functional NeuroImages (Cox 1996) and FMRIB Software Library suites (Smith et al. 2004). Skull-stripping was done using AFNI’s 3dskullstrip for the structural images and AFNI’s 3dautomask was used for the functional images. The functional images were then slice-time and head motion corrected registering each functional volume to the average functional volume using FSL’s FLIRT. For group comparisons, images were standardized to the UNC 0-1-2 neonate atlas using FSL’s nonlinear registration tool (FNIRT). In order to reduce motion artifacts, six rigid-body motion parameters were modeled as nuisance variables and removed with regression from all analysis. Motion for each time point, which is defined as root mean square of the sum displacement (RMSD) of all six translational and rotational axes was determined for each participant and used as a covariate in all group-level analyses. For any instance of RMSD >0.5mm, the time point was censored; and only participants with ≥80% (≥192) time points remaining were included. No significant group differences were found for total number of volumes scrubbed per group; an average of 7.12 volumes were removed for HR group (11 participants) and 6.76 for LR group (8 participants) (*t*(50)=.06, *p*=.95). In order to isolate spontaneous low-frequency BOLD fluctuations (Cordes et al. 2001), rs-fcMRI time series were bandpass filtered (.008 < *f* < .08 Hz), using a second-order Butterworth filter, which was also applied to all nuisance regressors described below. Images were spatially smoothed to a Gaussian full width at half-maximum (FWHM) of 5mm, using AFNI’s 3dBlurToFWHM.

Partial correlation analyses were performed in AFNI to obtain the unique connectivity pattern between each cortical ROI and thalamus bilaterally. The average BOLD time series was extracted from each cortical ROI and field of view (FOV) was restricted to bilateral thalamus. Partial correlations were computed between each bilateral cortical ROI and each voxel within bilateral thalamus, eliminating the shared variance by regressing out the time series from the other cortical ROIs. Two-sample, two-tailed t-tests were performed for between-group comparisons and all statistical maps were adjusted for multiple comparisons to a corrected *p*<.05 using AFNI’s 3dClustSim (https://afni.nimh.nih.gov/pub/dist/doc/program_help/3dClustSim.html). For all resulting significant clusters, parameter estimates of connectivity strength were extracted in AFNI and then correlated with behavioral measures.

#### DTI Data Processing

The diffusion weighted imaging data were checked for data quality using DTIPrep (Liu et al. 2010), which automatically detects and tracks bad gradient diffusion weighted images. Volumes were also visually inspected, and if the presence of residual artifacts was detected, these were then removed. Datasets including less than 23 (72%) gradient diffusion-weighted images were excluded due to low signal-to-noise ratio (Wolff et al. 2012). Thirteen datasets did not meet this threshold and were removed, resulting in a final sample of 34 infants (HR=19, LR=15) for these analyses. Groups did not differ significantly on average number of gradients remaining following quality control [Gradients remaining for HR: *M*=30.80, *SD*=1.90; Gradients remaining for LR: *M*=30.68, *SD*=1.67; *p*=.85]. DTI data were then preprocessed and analyzed using FSL’s FDT (FMRIB’s Diffusion Toolbox version 3.0 (Behrens et al. 2003; Behrens et al. 2007). FSL’s FDT includes motion correction, eddy current correction, and brain extraction. DTIFit was used to fit the diffusion tensor model at each voxel for fiber tracking. FSL was used to preprocess the T2 structural scan that was used to register diffusion data. Lastly, the DTI data were registered to a neonate template brain in standard space (Shi et al. 2011) using a 12-parameter affine transformation.

Probabilistic fiber tracking was performed using FDT to derive white matter tracts originating from each cortical seed and terminating at the thalamus. Unique exclusion masks were created for each target ROI (whole cortex minus specific target ROI) and the bilateral thalamus was selected as the termination mask. FSL’s BEDPOSTX was used to generate a Bayesian estimate of the probability distribution of different directions at each voxel. For each subject, all tractography outputs were first thresholded to exclude voxels with connectivity values less than 10,000 tracts. These tracts were then normalized to standard space and summed to create a normalized tract, which was thresholded to included voxels that belonged to a particular tract in at least 50% of the subjects within each group. Two additional HR participants were excluded during fiber tracking as their target tracts did not meet these thresholding standards resulting in a final sample of 17 HR and 15 LR infants. The diffusion tensor was calculated at each voxel and indices of white matter connectivity strength including FA, MD, RD, and AD were generated for tracts between each cortical ROI and thalamus. A one-way multivariate analysis of variance (MANOVA) was performed in SPSS software version 25.0 (IBM; Armonk, NY, USA) for between group comparisons for these DTI indices within each cortical ROI-thalamic tract. Pearson’s correlations were used to examine the relationship between DTI indices and behavioral measures.

#### Behavioral Data Analysis

The eye-tracking metrics and diagnostic measures used in our analyses were selected *a priori* to examine the relationship between imaging indices of thalamocortical connectivity and both early social functioning and risk for ASD. All the analyses correlating imaging indices and behavioral measures were Bonferroni-corrected to control for multiple comparisons.

For the ‘Charlie Brown’ eye-tracking task, our metrics of interest were percent fixation on any face during the depicted social interactions, and percent fixation to parts of the video clip of non-social perceptual salience (e.g., motion, high luminance, high saturation, edges). The areas of interest (faces, non-social salient features) were hand-traced similar to prior research with the same stimuli (Frank et al. 2014; Tsang *et al.* 2019) using software written in MATLAB (MathWorks, Inc; Natick, MA). We calculated the percentage of total fixations on faces and other salient perceptual features relative to the total duration of the video stimuli, respectively. Additionally, we calculated the percent of time looking at faces over time looking at salient perceptual features to examine preferential attention to social versus salient features within the visual field. Pearson’s correlations were conducted in SPSS between these eye-tracking metrics (collected at 6, 9, and 12 months of age) and both rs-fcMRI and DTI indices of connectivity strength for regions where significant group differences were observed. In the HR group only, these functional and structural connectivity indices were also correlated with the total scores on the AOSI, SRS-P, and ADOS-2. Spearman’s correlations were conducted in SPSS between the imaging indices and these diagnostic measures since they were not normally distributed.

## RESULTS

### Thalamocortical Connectivity

Rs-fcMRI results (Figure 1B) between the six cortical ROIs and bilateral thalamus indicated that in both LR and HR groups (Figure 1B; top panel, middle panel) parietal and occipital cortices were largely connected with the pulvinar, motor cortices were connected with the anterior ventral and mediodorsal nuclei, and somatosensory cortices were connected with the posterior ventral nuclei; temporal cortices showed robust connectivity with most thalamic nuclei at this age across both groups. Prefrontal cortices showed positive connectivity with anterior/mediodorsal nuclei of the thalamus in the LR group (Figure 1B; top panel); in contrast, the HR group displayed negative connectivity between prefrontal cortices and the anterior and mediodorsal (extending to lateral dorsal) nuclei of the thalamus (Figure 1B; middle panel). Direct between-group comparisons confirmed these differences to be statistically significant (Figure 1B; bottom panel), with prefrontal-thalamic networks showing significant underconnectivity in the HR group compared to LR group. Additionally, between-group comparisons also revealed that the HR group showed overconnectivity in thalamic-occipital and thalamic-motor networks as compared to the LR group (Figure 1B; bottom panel).

Given that the prefrontal cortex comprises a relatively large region of interest, we further examined which specific functional subregions of the prefrontal cortex might be driving the thalamic-prefrontal underconnectivity observed in the HR group. For this reverse functional connectivity analysis, the average BOLD time series was extracted from the thalamic cluster showing significant underconnectivity between the prefrontal cortical ROI and bilateral thalamus and then correlated with each voxel within the prefrontal ROI. Results from this analysis (Figure 1C) indicated that the thalamic-prefrontal underconnectivity observed in the HR group was driven more specifically by underconnectivity with the right inferior frontal gyrus.

Next, we examined whether white matter structural connectivity was impacted within the thalamocortical networks where we observed atypical connectivity in our rs-fcMRI analyses. That is, we examined group differences in DTI indices within the thalamic-prefrontal, thalamic-occipital, and thalamic-motor white matter tracts. Results from the MANOVA analyses run with all DTI indices for each thalamocortical tract revealed that, for the thalamic-occipital tract, all diffusivity indices - namely, MD (*p*=.03), RD (*p*=.02), AD (*p*=.05) - were significantly higher in the HR group compared to the LR group (Figure 2), though FA values did not differ significantly between the two groups (*p*=0.41). No other significant group differences were observed across any DTI indices for the other thalamocortical tracts.

**Figure 2.**
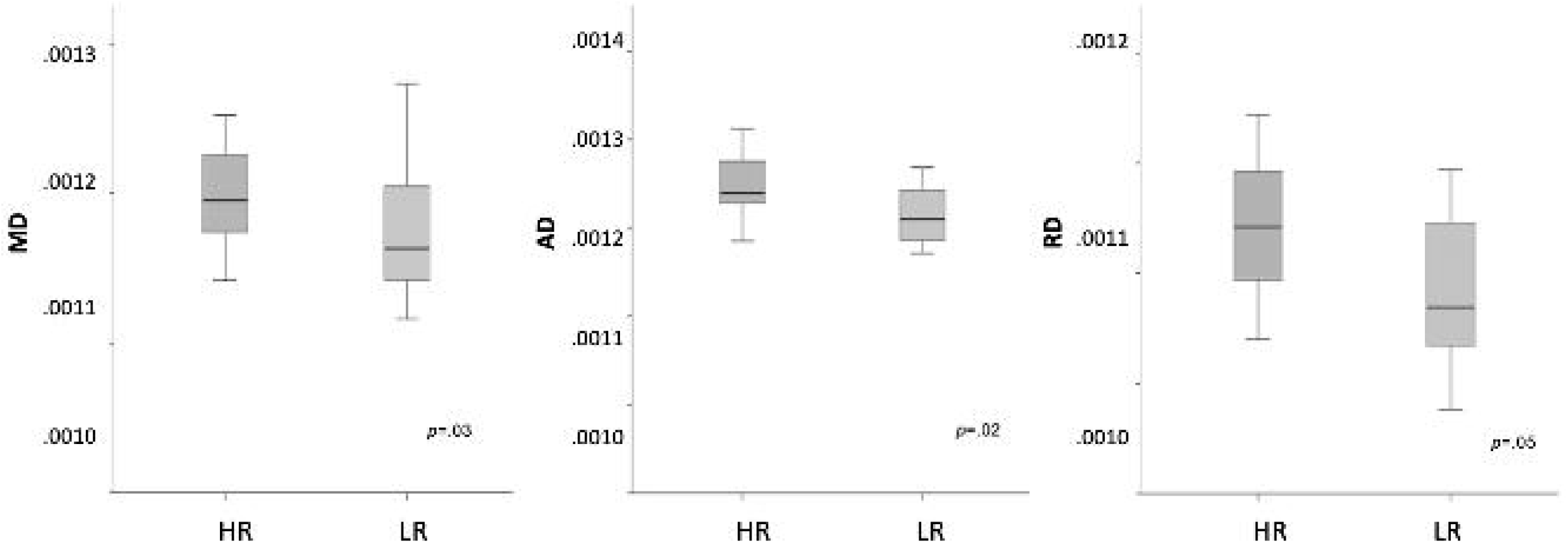
Diffusion tensor imaging thalamocortical connectivity results for 6-week old HR infants compared to their LR peers. Between-group results indicate that DTI diffusivity indices (MD, AD, RD) were significantly higher for HR group compared to LR group for thalamic-occipital tracts.

#### Brain-Behavior Relationships

Correlations were conducted between eye-tracking metrics and brain-based indices of functional and structural connectivity extracted from regions showing significant between-group differences (i.e., rs-fcMRI connectivity strength for thalamic-prefrontal, thalamic-occipital, and thalamic-motor networks; DTI diffusivity indices for thalamic-occipital network). In the LR group, greater thalamic-prefrontal rs-fcMRI connectivity strength was associated with higher percent time spent looking at faces (*r*=−.69, *p*=.002; Figure 3A), as well as greater preferences for faces (*r*=.63, *p*=.005; Figure 3B) over other perceptually salient features at age 9 months; similar patterns of correlations were observed in this group at the 6 and 12-month timepoints but these did not survive correction for multiple comparisons. For the HR group, higher thalamic-occipital RD (*r*=.52, *p*=.01; Figure 3C) was significantly correlated with greater preference for non-social perceptually salient features (vs. faces) at age 9 months; again, similar correlational patterns were also observed for this HR group at 6 and 12 months of age but these did not survive correction for multiple comparisons. No other significant correlations between brain-based measures and eye-tracking metrics survived correction for multiple comparisons in either group.

**Figure 3.**
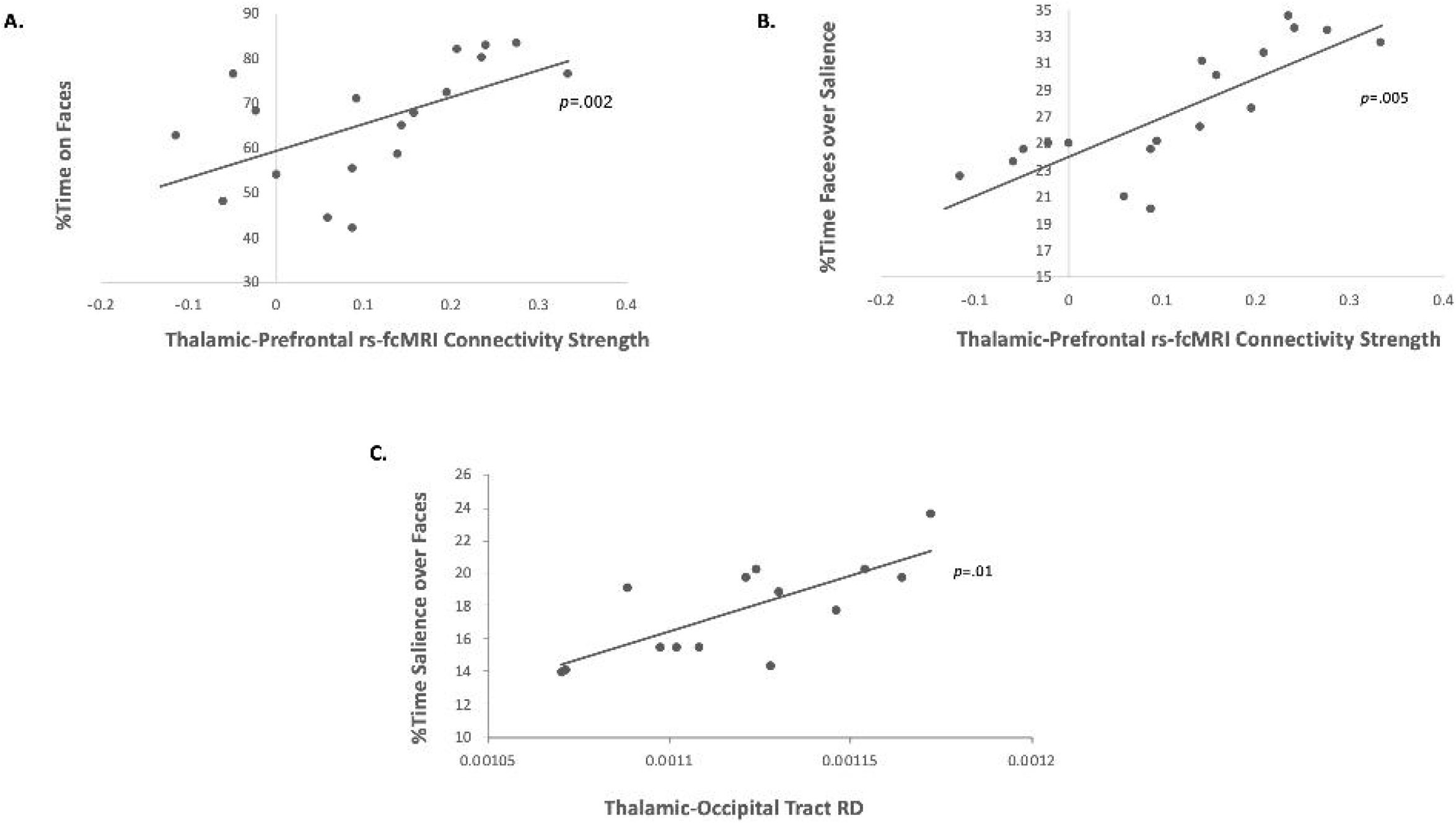
Correlations with eye-tracking indices on the “Charlie Brown” paradigm at age 9 months **(A.)** Correlation between percent time spent on faces and thalamic-prefrontal rs-fcMRI connectivity strength in the LR group. **(B.)** Correlation between percent time spent on faces versus other salient perceptual features and thalamic-prefrontal rs-fcMRI connectivity strength in the LR group. **(C.)** Correlation between percent time spent on salient perceptual features versus faces and thalamic-occipital tract RD in the HR group.

Lastly, for the HR group only, we examined the relationship between the same indices of functional and structural connectivity with caregiver ratings and clinician-administered measures of ASD risk. Results showed that higher thalamic-occipital rs-fcMRI connectivity strength was associated with higher scores on the AOSI obtained at age 12 months (*r*=.44, *p*=.03; Figure 4A) as well as higher scores on the SRS-P obtained at age 24 months (*r*=.58, *p*=.02; Figure 4B) and higher comparison scores on the ADOS-2 obtained at age 36 months (*r*=.52, *p*=.02; Figure 4C). No other significant correlations were observed between connectivity indices and both caregiver ratings and clinician-administered measures.

**Figure 4.**
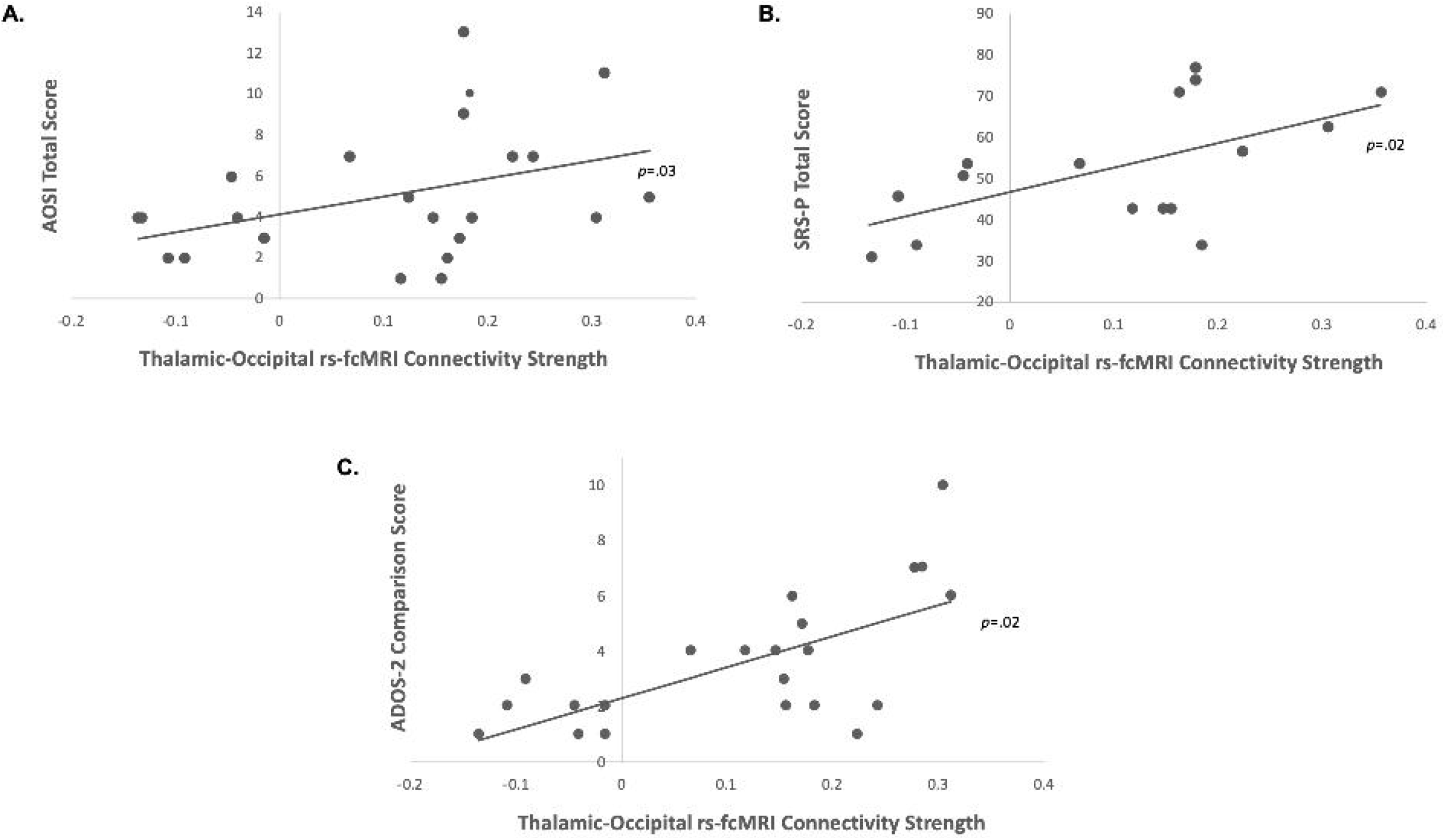
**(A.)** Correlations between rs-fcMRI connectivity strength for thalamic-occipital network and AOSI total raw score at 12 months for the HR group. **(B.)** Correlations between rs-fcMRI connectivity strength for thalamic-occipital network and SRS-P total T-score at 24 months for the HR group. **(C.)** Correlations between rs-fcMRI connectivity strength for thalamic-occipital network and ADOS-2 comparison score at 36 months for the HR group.

## DISCUSSION

This is the first multimodal study to examine both the functional and structural thalamocortical connectivity in infants at high (HR) and low (LR) familial risk for ASD as early as 6-weeks post birth. Overall, our findings indicate that thalamocortical connectivity patterns in LR infants were similar to what observed in prior studies in older neurotypical individuals (Zhang *et al.* 2008; Fair *et al.* 2010; Zhang *et al.* 2010; Nair *et al.* 2013; Nair *et al.* 2015). In contrast, connectivity of thalamocortical networks in the HR group was atypical with significant functional underconnectivity observed in thalamic-prefrontal networks, accompanied by functional overconnectivity in thalamic-motor and thalamic-occipital networks. Structural connectivity of thalamocortical networks was also impacted with higher white matter diffusivity observed in thalamic-occipital tracts in HR infants as compared to LR infants. Interestingly, indices of both functional and structural thalamocortical connectivity at six weeks of age were predictive of later social development and risk for ASD. More specifically, stronger thalamic-prefrontal functional connectivity in LR infants was associated with greater attention to faces (vs. non-social but salient visual information) as assessed with eye-tracking from 6 to 12 months of age; in contrast, aberrant connectivity in thalamic-occipital networks in HR infants was associated with visual preference for non-social information on the eye-tracking task as well as increased ASD symptomatology from 12 to 36 months, as indexed by both caregivers’ report and clinician administered measures.

### Aberrant Connectivity Patterns

In LR infants, thalamocortical connectivity patterns generally mirrored those reported in prior studies in older neurotypical individuals (Zhang *et al.* 2008; Fair *et al.* 2010; Zhang *et al.* 2010; Nair *et al.* 2013; Nair *et al.* 2015) whereby prefrontal cortices show strongest connections with anterior and medial nuclei of the thalamus, motor cortices with ventral anterior and lateral nuclei, somatosensory cortices with ventral posterior nuclei, parietal and occipital cortices with lateral pulvinar, and temporal cortices with medial pulvinar. While it is hard to accurately identify thalamic nuclei in such young infants, it appears that our findings in LR infants closely match this normative pattern observed in adolescents and adults, except for temporal cortices which showed significant functional connectivity with almost all thalamic nuclei. Our results are also consistent with prior studies on thalamocortical functional connectivity in full-term new-born infants (Alcauter et al. 2014; Toulmin et al. 2015; Cai et al. 2017) demonstrating thalamocortical network organization similar to what observed in older individuals, despite the use of slightly different regions of interest. Interestingly, pre-term infants (gestational age=24-32 weeks) instead showed thalamic underconnectivity with prefrontal regions and overconnectivity with post-central gyri compared to full-term infants (Toulmin *et al.* 2015). Thus, the patterns of thalamocortical connectivity observed in HR infants in the current study are similar to those seen in pre-term infants in that thalamic-prefrontal networks were underconnected, whereas thalamic connectivity with primary sensory regions were overconnected.

This pattern of reduced connectivity between thalamus and supramodal cortical regions (prefrontal), and increased connectivity between thalamus and primary cortical regions (visuo-motor) in HR infants in this study also aligns well with the altered thalamocortical connectivity previously observed in youth with ASD (Nair *et al.* 2013; Nair *et al.* 2015). In these prior thalamocortical studies, functional underconnectivity was reported with prefrontal, parieto-occipital, motor, and somatosensory cortices, along with overconnectivity with temporal and limbic cortices. However, DTI indices in the underlying thalamocortical white matters tracts in these prior studies were found to be mostly impacted in frontal, temporal, and parietal regions, in contrast to our current findings of impacted white matter connectivity in thalamic-occipital tracts in HR infants. Interestingly, a study by (Fair *et al.* 2010) examining the typical maturational trajectories of these thalamocortical networks from childhood to adulthood suggested an increase in connectivity within thalamic-prefrontal, thalamic-motor, and thalamic-somatosensory networks with age, whereas thalamic-temporal connectivity typically appeared to weaken with increasing age. Hence, our prior findings on thalamocortical connectivity in youth with ASD as well as the current findings in HR infants suggest that both of these groups may be on a delayed and/or differential maturational trajectory for subcortical-cortical connectivity compared to their typically-developing peers. Notably, however, thalamic-occipital networks showed a quadratic developmental trajectory in healthy controls with connectivity peaking in adolescent years and weakening with age (Fair *et al.* 2010); thus, the early thalamic-occipital overconnectivity in HR infants and subsequent underconnectivity in ASD youth observed in the current and prior studies suggest a more complicated disruption in the maturation of this network that may be linked to distinct behavioral symptoms at each developmental period.

### Associations with Social Attention and Risk for ASD

The atypicalities in early brain connectivity we observed in the present study, and their relationships with later eye-tracking and behavioral measures, suggest disruptions in specific thalamocortical networks implicated in social attention and engagement in HR infants. In LR infants, thalamic-prefrontal functional connectivity was associated with greater attention to social stimuli such as faces in the first year of life. In contrast, this relationship was not observed in HR infants who instead showed thalamic underconnectivity with prefrontal cortex, particularly the right inferior frontal gyrus. Notably, this region has been implicated in endogenous as well as exogenous attentional shifts, (Peelen et al. 2004; de Fockert and Theeuwes 2012; Caruana et al. 2015; Koike et al. 2016), as well as social cognition processes (Hartwigsen et al., 2019; Sato et al., 2019; Sokolow et al., 2018). Accordingly, thalamic underconnectivity with this region may underlie later difficulties in orienting attention to socially relevant information in HR infants. Interestingly, this aberrant functional and structural connectivity in thalamic-occipital networks in HR infants was associated with an atypical visual preference for non-social information as well as increased ASD symptoms as observed by caregivers and clinicians from 12 to 36 months. As thalamic-occipital networks are involved in modulation and orientation of visual engagement (Carrera and Bogousslavsky 2006; Arend et al. 2008; Snow et al. 2009; Li et al. 2018), early disruptions in the neural development of these networks may contribute to atypicalities in the processing of visual information. the group differences we observed in the relationships between thalamocortical connectivity and preferential viewing patterns during the first year of life suggest greater salience of non-social visual features, vs. socially-relevant stimuli, for HR infants as well as a possible very early biomarker for ASD risk.

A number of eye-tracking studies in HR infants have previously highlighted that disrupted attention to and visual processing of social information may be a precursor to eventual diagnosis of ASD. Collectively, this body of work has demonstrated diminished attention to faces (Elsabbagh *et al.* 2012; Chawarska *et al.* 2013; Jones and Klin 2013; Chawarska et al. 2016; Constantino et al. 2017) and biological motion (Klin et al. 2009; Klin, Klaiman, et al. 2015) in infants and toddlers who eventually receive an ASD diagnosis. Importantly, neuroimaging findings have also shown that network atypicalities in frontal language regions and white matter tracts involved in low-level sensory processing in HR infants as young as 6 months of age are predictive of an eventual ASD diagnosis (Lewis *et al.* 2014; Lewis *et al.* 2017). The current study extends these prior findings and highlights the role of early thalamic-prefrontal connectivity in scaffolding normative social attention as well as the involvement of altered thalamic-occipital connectivity for both atypical visual processing of social information and increased risk for ASD. Taken together, the present findings suggest that atypical subcortical connectivity with prefrontal and occipital regions might provide one of the earliest indicators of aberrant brain development in HR infants.

Atypical social attention has also been demonstrated in prior research with older youth with ASD (Klin et al. 2002; Caron et al. 2006; McPartland et al. 2011; Samson et al. 2012; Guillon et al. 2014; Chita-Tegmark 2016), together with findings of reduced functional connectivity in social cognition networks involving prefrontal, limbic, and posterior temporal brain regions in ASD individuals (Pelphrey et al. 2011; Gotts et al. 2012; von dem Hagen *et al.* 2013; Cheng et al. 2015; Linke *et al.* 2018; Muller and Fishman 2018; Delbruck et al. 2019; Maximo and Kana 2019; Odriozola et al. 2019; Sato and Uono 2019). Importantly, key ASD symptomatology including atypical social attention and repetitive stereotyped behaviors have also been previously linked to thalamocortical network disruptions in youth with ASD (Nair *et al.* 2013; Nair *et al.* 2015). Our current findings thus suggest that very early disruptions in key thalamocortical circuits involved in social-visual engagement – such as thalamic-prefrontal and thalamic-occipital networks – may underlie risk for atypical social development in HR infants, preceding and perhaps contributing to the overt manifestation of overt ASD symptoms later in development. Further longitudinal investigations in a larger sample are clearly needed to determine the predictive value of early atypical thalamocortical connectivity in identifying HR infants who will later receive a diagnosis of ASD, vs. those who will show typical development or other suboptimal outcomes.

### Limitations and Conclusions

This study has several limitations. As mentioned above, the overall limited sample size only allowed us to identify atypicalities associated with increased risk for ASD, as opposed to those predictive of a subsequent ASD diagnosis. The sample size for our structural connectivity analyses was further reduced, relative to the sample used in our analyses of functional connectivity, as infants often woke up more before or during the DTI sequence which was acquired in the MRI session. Reduced power for these analyses could partly explain the limited between-group differences observed in structural connectivity (after correction for multiple comparisons). Additionally, several participants had missing or unusable eye-tracking data across the three different timepoints (6, 9, and 12 months of age); this may have impacted the outcomes of our correlational analyses with eye-tracking metrics, resulting in trend-level associations with atypical connectivity indices for some of the timepoints that did not survive correction for multiple comparisons.

To the best of our knowledge, this is the first study to examine thalamocortical functional and structural connectivity in new-born infant siblings at high familial risk for ASD. Our results indicate disruptions in functional and structural thalamocortical connectivity occurring very early in life in infants at high familial risk for ASD and implicate early thalamocortical connectivity in both typical and atypical social development. Our findings further our understanding of the early neural mechanisms underlying risk for the ASD and have implications for identifying earlier time frames for screening HR infants as well as informing targeted interventions that may help steer brain development toward more normative pathways.

## ACKNOWLEDGMENTS

This work was supported by the National Institute of Child Health and Human Development (NICHD P50 HD055784), and the Autism Science Foundation Postdoctoral Fellowship 16-004 (author AN). We are also grateful for the generous support from the Brain Mapping Medical Research Organization, Brain Mapping Support Foundation, Pierson-Lovelace Foundation, The Ahmanson Foundation, Capital Group Companies Charitable Foundation, William M. and Linda R. Dietel Philanthropic Fund, and Northstar Fund. Research reported in this publication was also partially supported by the National Center for Research Resources and by the Office of the Director of the National Institutes of Health under award numbers C06RR012169, C06RR015431 and S10OD011939. The content is solely the responsibility of the authors and does not necessarily represent the official views of the National Institutes of Health.

Human subjects’ oversight and approval was provided by UCLA’s Institutional Review Board. Special thanks to the participants and their families, and staff and volunteers who contributed toward imaging data collection.

## CONFLICT OF INTERESTS

The authors declare that they have no competing interests.

## Notes

### Competing Interest Statement

The authors have declared no competing interest.

