## Supplementary Material for "Altered Thalamocortical Connectivity in Six-Week Old Infants at High Familial Risk for Autism Spectrum Disorder"

Supplementary Table 1. UNC Infant 0-1-2 neonate atlas anatomical parcellations used to create cortical ROIs

| **Prefrontal** | **Parietal** | **Occipital** | **Motor** | **Somatosensory** | **Temporal** |
| --- | --- | --- | --- | --- | --- |
| SFGdor-L | PCG-L | CAL-L | SMA-L | PoCG-L | HIP-L |
| SFGdor-R | PCG-R | CAL-R | SMA-R | PoCG-R | HIP-R |
| ORBsup-L | SPG-L | CUN-L | PreCG-L | PCL-L | PHG-L |
| ORBsup-R | SPG-R | CUN-R | PreCG-R | PCL-R | PHG-R |
| MFG-L | IPL-L | LING-L |  |  | AMYG-L |
| MFG-R | IPL-R | LING-R |  |  | AMYG-R |
| ORBmid-L | SMG-L | SOG-L |  |  | FFG-L |
| ORBmid-R | SMG-R | SOG-R |  |  | FFG-R |
| IFGoperc-R | ANG-L | MOG-L |  |  | HES-L |
| IFGoperc-R | ANG-R | MOG-R |  |  | HES-R |
| IFGtriang-L | PCUN-L | IOG-L |  |  | STG-L |
| IFGtriang-R | PCUN-R | IOG-R |  |  | STG-R |
| ORBinf-L |  |  |  |  | TPOsup-L |
| ORBinf-R |  |  |  |  | TPOsup-R |
| SFGmed-L |  |  |  |  | MTG-L |
| SFGmed-R |  |  |  |  | MTG-R |
| ORBmed-L |  |  |  |  | TPOmid-L |
| ORBmed-R |  |  |  |  | TPOmid-R |
| REC-L |  |  |  |  | ITG-L |
| REC-R |  |  |  |  | ITG-R |
| ACG-L |  |  |  |  |  |
| ACG-R |  |  |  |  |  |
| MCG-L |  |  |  |  |  |
| MCG-R |  |  |  |  |  |

Supplementary Figure 1. “Charlie Brown” eye-tracking paradigm

*p <.05 corr.*

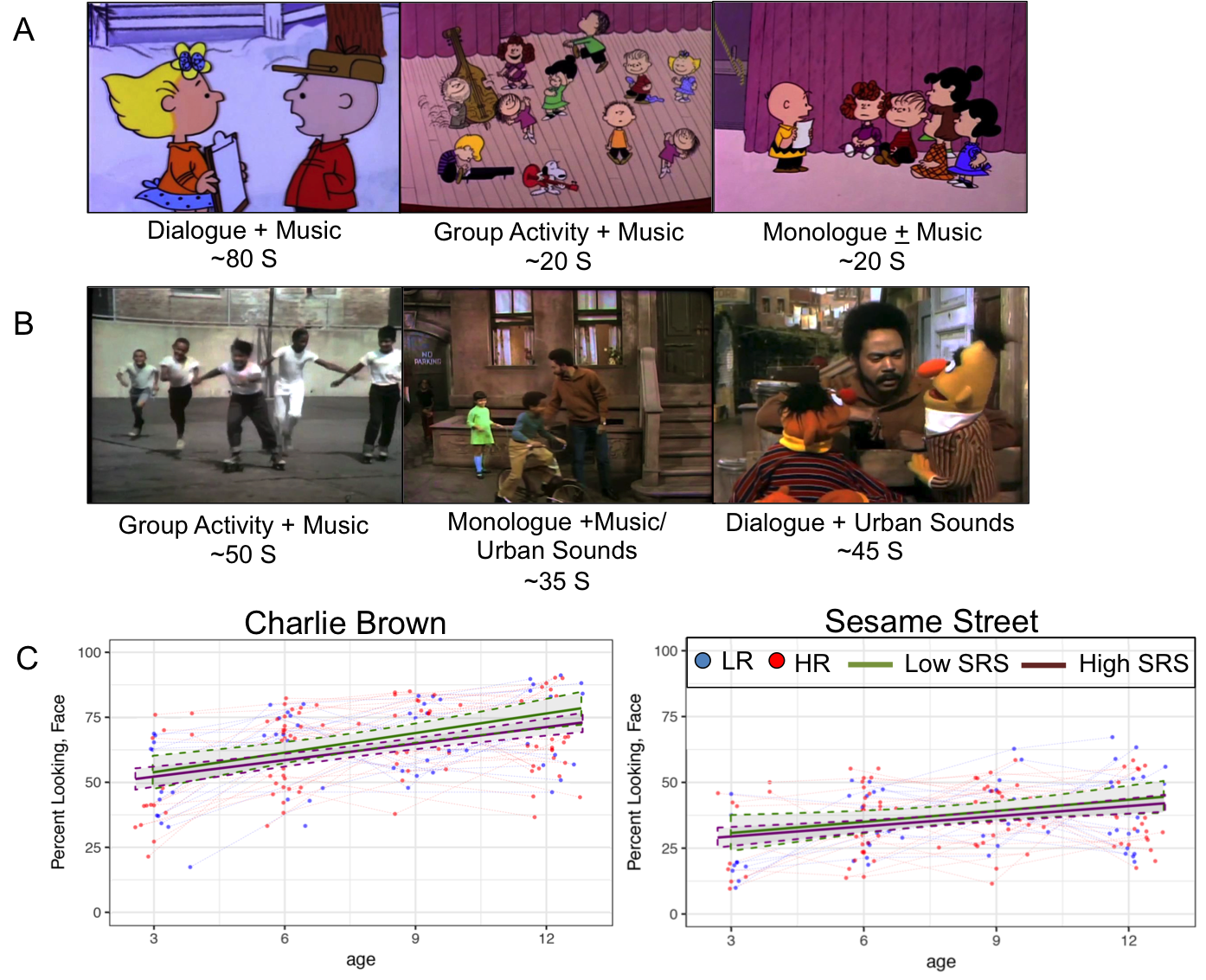
